# A multistage *Plasmodium* CRL4^WIG1^ ubiquitin ligase is critical for the formation of functional microtubule organisation centres in microgametocytes

**DOI:** 10.1101/2023.07.19.549332

**Authors:** Ravish Rashpa, Cameron Smith, Katerina Artavanis-Tsakonas, Mathieu Brochet

## Abstract

Malaria is a mosquito-borne infectious disease caused by unicellular eukaryotic parasites of the *Plasmodium* genus. Protein ubiquitination by E3 ligases is a critical post-translational modification required for various cellular processes during the lifecycle of *Plasmodium* parasites. However, little is known about the repertoire and function of these enzymes in *Plasmodium.* Here we show that *Plasmodium* expresses a conserved cullin RING E3 ligase (CRL) complex that is functionally related to the eukaryotic CRL4. In *P. falciparum* asexual blood stages, a cullin-4 scaffold interacts with the RING protein RBX1, the adaptor protein DDB1 and a set of putative receptor proteins that may determine substrate specificity for ubiquitination. These receptor proteins contain WD40-repeat domains and include <u>W</u>D-repeat protein Important for <u>G</u>ametogenesis 1 (WIG1). This CRL4-related complex is also expressed in *P. berghei* gametocytes, with WIG1 being the only putative receptor detected in both schizont and gametocyte stages. While WIG1 is not required for the proliferation of *P. berghei* asexual blood stages, its disruption leads to a complete block in microgamete formation. Proteomic analyses indicate that *WIG1* disruption alters proteostasis of ciliary proteins and components of the DNA replication machinery during gametocytogenesis. Further analysis by ultrastructure expansion microscopy (U-ExM) indicates that WIG1-dependent depletion of ciliary proteins is associated with impaired formation of the microtubule organisation centres that coordinate mitosis with axoneme formation and altered DNA replication during microgametogenesis. This work identifies a CRL4-related ubiquitin ligase in *Plasmodium* that is critical for the transmission of malaria parasites by regulating proteostasis of ciliary and DNA replication proteins.

## Introduction

Malaria remains a major global health problem with 247 million people infected and 0.6 million deaths in 2021 [1]. The lifecycle of *Plasmodium* parasites alternates between mosquito and vertebrate hosts. Clinical symptoms of malaria are linked to the dividing asexual blood stages, whereas transmission to the mosquito is solely initiated by sexually differentiated gametocytes. Differentiation from asexually replicating stages into gametocytes takes place inside host erythrocytes. Following a period of maturation, micro– and macrogametocytes are ready to initiate transmission when ingested by a mosquito during a bloodmeal. Gametocytes resume their development in response to environmental conditions encountered in the mosquito midgut. These include a small mosquito molecule, xanthurenic acid (XA), a rise in extracellular pH, and a drop in temperature. Upon activation, gamete egress from the host erythrocyte relies on the release of secretory vesicles called osmiophilic bodies [2]. The macrogamete is rapidly available for fertilisation following activation of translationally repressed mRNAs [3, 4]. In comparison, the microgametocyte undergoes a more dramatic developmental process by completing three rounds of genome replication and closed mitosis, by assembling eight axonemes, and finally by producing eight haploid microgametes in a process called exflagellation, all in less than 15 minutes.

Microgametogenesis relies on the synchronisation of mitosis and axoneme assembly by bipartite microtubule organization centers (MTOC). In non-activated microgametocytes, a single bipartite MTOC lies on the cytoplasmic face of a nuclear pore that is physically linked to an intranuclear body in the nuclear face of the same pore [9]. At this stage, centrin, γ-tubulin, SAS4-HA and SAS6-HA are present in the amorphous MTOC but are not incorporated into a structure resembling known MTOC such as centrosome, basal body, or spindle pole body. Upon activation of gametogenesis, the MTOC gives rise to two tetrads of four basal bodies from the amorphous MTOC through the dynamic relocalisation of SAS4, SAS6 and centrin. On the nuclear side, the intranuclear bodies differentiate into spindle poles that coordinate concomitant mitotic events [5]. After 1-2 min, the first mitotic spindle is visible, and four basal bodies with growing axonemes that display the canonical 9+2 microtubule architecture are found at each extremity of the mitotic spindle. Each of the eight axoneme polymerise in the cytoplasm in absence of intra-flagellar transport [6, 7]. Three to five minutes later mitosis II occurs, during which a pair of basal bodies are found at each end of two mitotic spindles. By 6-8 min after initiation, the four spindles of mitosis III are visible and a single basal body is found attached to each of the eight spindle poles. The processes of chromatin condensation begins at the end of mitosis III, when each of the eight short spindles anchors one haploid set of 14 chromosomes to its respective spindle pole. At the onset of exflagellation, the axonemes become motile and incorporate a haploid genome into the forming flagellated gamete [7].

This coordination between genome replication, closed mitosis and axoneme assembly requires tight regulation. The role of protein phosphorylation by kinases has been shown to be crucial for the formation of microgametes [8, 9]. This includes a kinase related to cyclin-dependent kinases [10], a mitogen-activated protein kinase [11, 12], an aurora kinase [13] or plant-like calcium-dependent protein kinases [14–18]. More recently, ubiquitination by ubiquitin ligases has emerged as a critical protein post-translational modification in the regulation of *Plasmodium* gametogenesis. A first snapshot of the gametocyte ubiquitome revealed 2,183 ubiquitinated lysine residues mapping on 519 proteins [19]. Ubiquitination is mediated by various E3 ubiquitin ligases, grouped by their composite protein domains: HECT, U-box, PHD-finger, and RING-finger domains. So far, proteins belonging to two complexes of the RING-finger family have been shown to play important roles during male gametogenesis. The anaphase-promoting complex/cyclosome (APC/C) is a ligase that targets mitotic proteins involved in the metaphase-to-anaphase transition in other eukaryotes [20]. Genome mining identified only four APC/C subunits in *Plasmodium*: a single CDC20/CDH1 homolog, APC3, APC10, and APC11 [21–23]. CDC20 and APC3 were shown to be critical for exflagellation with proposed roles for mitotic spindle formation but not DNA replication nor axoneme assembly [21, 22]. More recently, another multisubunit E3 ubiquitin ligase complex, the SKP1/Cullin-1/FBXO1 complex (CRL1 or SCF^FBXO1^), was shown the be crucial for multiple developmental stages during the *Plasmodium* lifecycle. This includes microgametogenesis during which, deletion or depletion of FBXO1 led to defects in the MTOC segregation and egress from the host red blood cell [19].

The CRL1 complex belongs to the class of cullin-RING ligases (CRLs), which are the largest family of ubiquitin E3 ligases, with over 400 members [24]. Mammalian cells express eight classes of CRL complexes, each containing at least four core components: one of eight cullin isoforms serving as a common backbone scaffold protein, one of two RING finger protein (Rbx1 or Rbx2) that binds to an E2 ubiquitin conjugating enzyme, a substrate receptor that recognize the target protein, and adaptor subunits that serve as links between the cullin scaffold and substrate receptors. Cells contain a multitude of substrate receptors, which are critical for the specificity of a given CRL for its target proteins. For example, in human, around 70 F-box protein receptors have been identified for CRL1. Despite the diversity of receptor proteins, they typically display protein–protein interaction domains such as tryptophan-aspartic acid 40 (WD40), leucine rich repeat (LRR), Src Homology 2 (SH2), or Kelch domains that recruit specific substrates for ubiquitination.

In addition to CUL1, *Plasmodium* encodes a second cullin annotated as cullin-2 (CUL4-PF3D7_0629800) that, compared to human cullins, shares maximum sequence identity with Cullin-4B. It was recently shown it forms a CRL4-related complex in *P. falciparum* asexual blood stages and is required for both intra erythrocytic proliferation and gametocytogenesis [25]. To keep in line with the nomenclature, we here propose to rename this protein CUL4. Here we confirm that CUL4 forms a CRL4-related complex in *P. falciparum* asexual blood stages that consists of RBX1, the adaptor protein DDB1 and a distinct set of putative receptor proteins containing WD40 repeat domains. We further show that this CRL4-related complex is expressed in *P. berghei* gametocytes with at least one putative receptor WD protein expressed at both stages. While this WD protein is not required for the proliferation of asexual blood stages, its disruption leads to a complete block in microgamete formation. We thus decided to name this protein <u>W</u>D repeat protein Important for <u>G</u>ametogenesis 1 (WIG1). Proteomic analyses indicate that *WIG1* disruption alters proteostasis of ciliary proteins and components of the DNA replication machinery during gametocytogenesis. Further analysis by ultrastructure expansion microscopy (U-ExM) indicates that WIG1-dependent depletion of ciliary proteins is associated with altered formation of the microgametocyte MTOCs and defects in DNA replication.

## Results

### CRL4-related complexes are expressed in *P. falciparum* schizonts

We first aimed at defining the protein partners of Cullin-4 (PfCUL4 – PF3D7_0629800) in *P. falciparum* schizonts. To do so, we generated a *P. falciparum* line expressing a HA-tagged PfCUL4 (Fig. S1A). Immunoblotting confirmed expression of the fusion protein at all stages during intraerythrocytic development with the expected size-dependent protein mobility (Fig. 1A). Immunofluorescence assays showed a cytoplasmic distribution of PfCUL4-HA with however no signal observed around hemozoin crystals suggesting its exclusion from the food vacuole (Fig. 1B).

**Figure 1.**
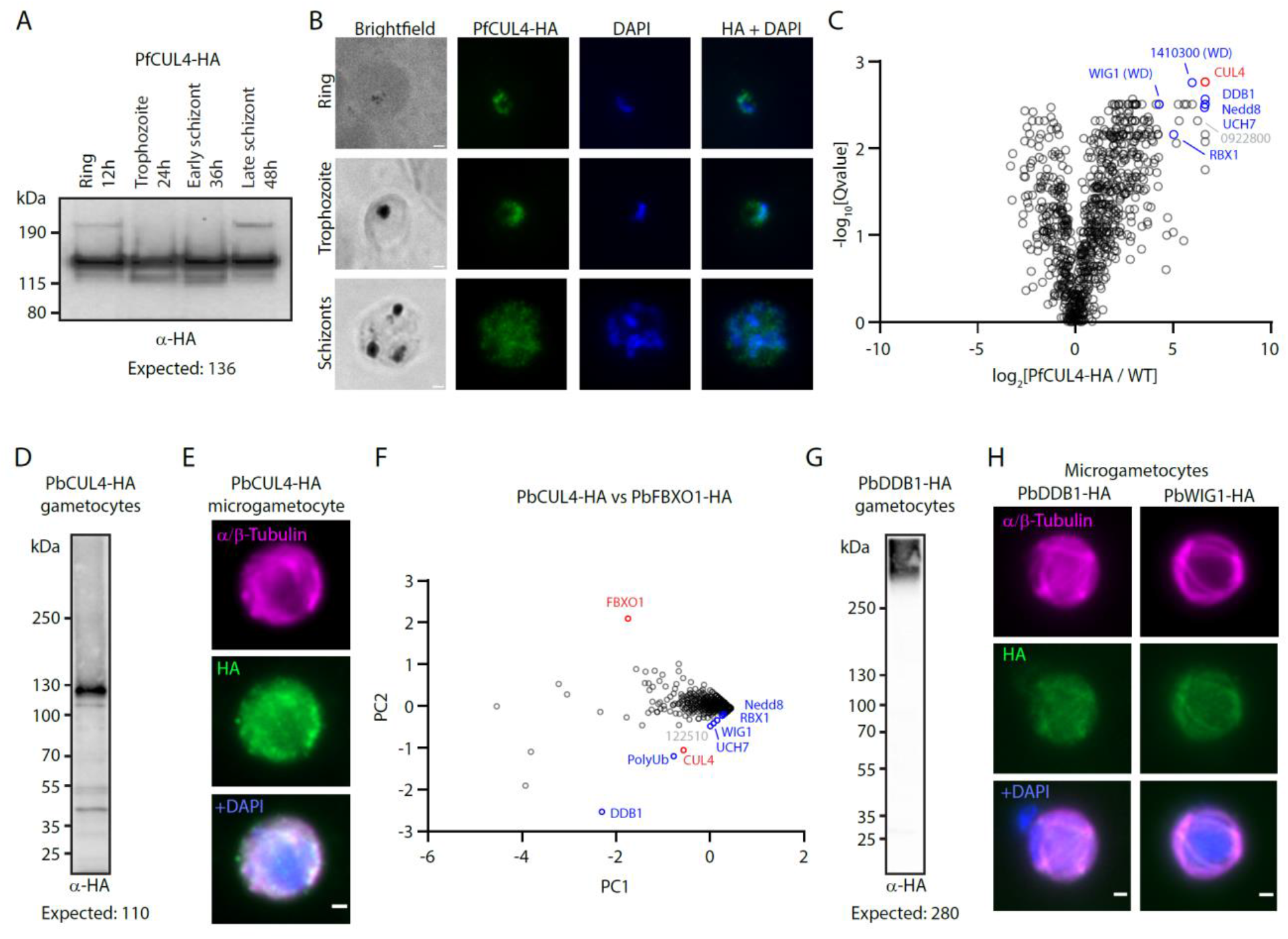
Expression of a CRL4^WIG1^ ubiquitin ligase complex in *P. falciparum* schizonts and *P. berghei* gametocytes. A. Western blot analysis of CUL4-HA expression in *P. falciparum* lysates collected at different time points during an intra erythrocytic cycle. **B.** Localisation of PfCUL4-HA (green) by widefield immunofluorescence in ring, trophozoite and schizont stages. DNA is stained with DAPI (blue). Scale bar = 1 µm. **C.** Volcano plot depicting relative abundance of proteins identified by semi-quantitative mass spectrometry in pull-downs from *P. falciparum* CUL4-HA compared to WT (control) parasites (n=3). Significance is expressed as log_10_ of the Q-value (y-axis). X-axis: enrichment of interaction partners in the *P. falciparum* CUL4-HA pull-down compared to controls. In red is the CUL4-HA bait. In blue and grey are components of the CRL4 complex together with a protein of unknown function also identified in reference [25], respectively. **D.** Western blot analysis of CUL4-HA expression in lysates of non-activated *P. berghei* gametocytes. **E.** Localisation of PbCUL4-HA (green) by widefield immunofluorescence in *P. berghei* gametocytes 8-10 minutes post-activation by XA. DNA is stained with DAPI (blue). Scale bar = 1 µm. F. Quantitative value (Normalized Total Spectra) for proteins co-purifying with PbCUL4-HA and PbFBXO1-HA (control) following immunoprecipitation, and displayed in first and second principal components (n = 3 biological replicates). Red denotes immunoprecipitated proteins and blue possible components of the CRL4 complex. **G.** Western blot analysis of DDB1-HA expression in lysates of non-activated *P. berghei* gametocytes. **H.** Localisation of PbDDB1-HA and PbWIG1-HA (green) by widefield immunofluorescence in *P. berghei* gametocytes 8-10 minutes post-activation by XA. DNA is stained with DAPI (blue). Scale bars = 1 µm.

To identify interacting proteins for PfCUL4, we then used affinity purification of PfCUL4-HA from synchronised *P. falciparum* schizonts followed by immunoprecipitation mass spectrometry (IP-LC-MS/MS) (Fig. 1C and Table S1). Supporting the notion of a CUL4-based CRL complex, the RING finger protein RBX1 (PF3D7_0319100) and Nedd8 (PF3D7_1313000) were enriched proteins co-purifying with PfCUL4-HA (Fig. 1c). Nedd8 shares 60% amino acid sequence identity to ubiquitin and is known to target cullins to activate their respective CRL complex. Interestingly, one of the most enriched protein besides PfCUL4 was a putative orthologue of DNA damage-binding protein 1 (DDB1 – PF3D7_0317700, currently annotated as Cleavage and Polyadenylation Specific Factor subunit A, CPSF_A), the main adaptor of CUL4-based CRL complexes in eukaryotes, further suggesting the conservation of a CUL4-based complex in *P. falciparum* asexual blood stages, as previously proposed [25]. Among the proteins significantly enriched in PfCRL4-HA immunoprecipitates were also two proteins containing WD domains (PF3D7_0321800 or WIG1, and PF3D7_1410300) that could represent possible substrate receptors of the *Plasmodium* CRL4-related complex. We also identified a ubiquitin carboxyl-terminal hydrolase (PfUCH7 – PF3D7_0726500) as significantly enriched in PfCUL4-HA immunoprecipitates. These significantly enriched proteins, together with a conserved *Plasmodium* protein of unknown function, PF3D7_0922800, were also recently identified in immunoprecipitates of PfCUL4-HA from asexual blood stages [25].

## The expression of a CRL4 complex is conserved in *P. berghei* gametocytes

As CUL4 and all its identified partners also share a transcription peak in sexual stages, we asked whether the CRL4-related complex was also conserved in gametocytes. To do so, we turned to the rodent malaria parasite *P. berghei* and generated line expressing a triple HA-tagged PbCUL4 (PBANKA_1128600 – Fig. S1B). Immunoblotting confirmed expression of the fusion protein in mature gametocytes with the expected mobility (Fig. 1D). No differential detection of the protein by western blot was observed during the ten first minutes following activation of gametocytes by XA (Fig. S1C). Immunofluorescence assays also showed a cytoplasmic distribution of CUL4-HA both in *P. berghei* schizonts (Fig. S1D) and gametocytes (Fig. 1E).

Affinity purification of PbCUL4-HA from 4 min activated *P. berghei* gametocytes also identified RBX1 (PBANKA_0806200), DDB1 (PBANKA_0807500), Nedd8 (PBANKA_1411400), and polyubiquitin (PolyUb – PBANKA_0610300) highlighting the conservation of a CRL4-based complex in multiple stages of the *Plasmodium* lifecycle (Fig. 1F and Table S2). Interestingly, the orthologues of the WD containing protein WIG1 (PBANKA_1216700) and the ubiquitin carboxyl-terminal hydrolase (PbUCH7 – PBANKA_0210600) were also enriched. To confirm these interactions, we generated *P. berghei* lines expressing HA-tagged alleles of PbDDB1 and PbWIG1 (Fig. S1B). While immunoblotting confirmed expression of DDB1-HA (Fig. 1G) in gametocytes, no signal by western blotting was detected for WIG1-HA despite in frame integration of the tag coding sequence (Fig. S1E). Immunofluorescence assays also indicated a cytoplasmic distribution of PbDDB1-HA and PbWIG1-HA in schizonts (Fig. S1D) and gametocytes (Fig. 1H). Affinity purification of PbDDB1-HA and PbWIG1-HA from 4 min activated *P. berghei* gametocytes followed by label-free semiquantitative mass spectrometry confirmed enrichment of PbCUL4, PbNedd8, PbRBX1, PbDDB1 and PbWIG1 compared with previous immunoprecipitations of the FBXO1 substrate receptor of the SCF1/CRL1 complex (Fig. S1F). In addition, another conserved *Plasmodium* protein of unknown function, PBANKA_1225100, was also enriched in PbCUL4-HA and PbDDB1-HA immunoprecipitates from *P. berghei* gametocytes.

Altogether these results highlight the conserved expression of a CUL4/RBX1/DDB1/WIG1 complex in both *P. falciparum* schizonts and *P. berghei* gametocytes.

## The putative CRL4 receptor WIG1 is essential for *P. berghei* male gamete formation

Downregulation of PfCUL4 previously pointed to requirement for CRL4 during both asexual replication and gametocytogenesis [25]. We then set out to further define the requirement of the CRL4^WIG1^ complex by focusing on WIG1. A *P*. *falciparum* global transposon mutagenesis project suggested PfWIG1 to be essential for asexual blood stages based on the absence of transposon insertion in the 9419 bp *WIG1* coding sequence [26]. We took advantage of the highly efficient PlasmoGEM vectors to attempt to delete the last 217 codons of the *WIG1* gene in *P. berghei* (Fig. S2). A WIG1-GD line was readily obtained and cloned suggesting no defect in the proliferation of asexual blood stages. Similarly, the WIG1-GD line produced microscopically normal female and male gametocytes, as assessed by Giemsa staining (Fig. 2A). However, upon activation by XA, no exflagellation events could be observed while, on average, 58% of WT male gametocytes led to active exflagellation centres (Fig. 2B).

**Figure 2.**
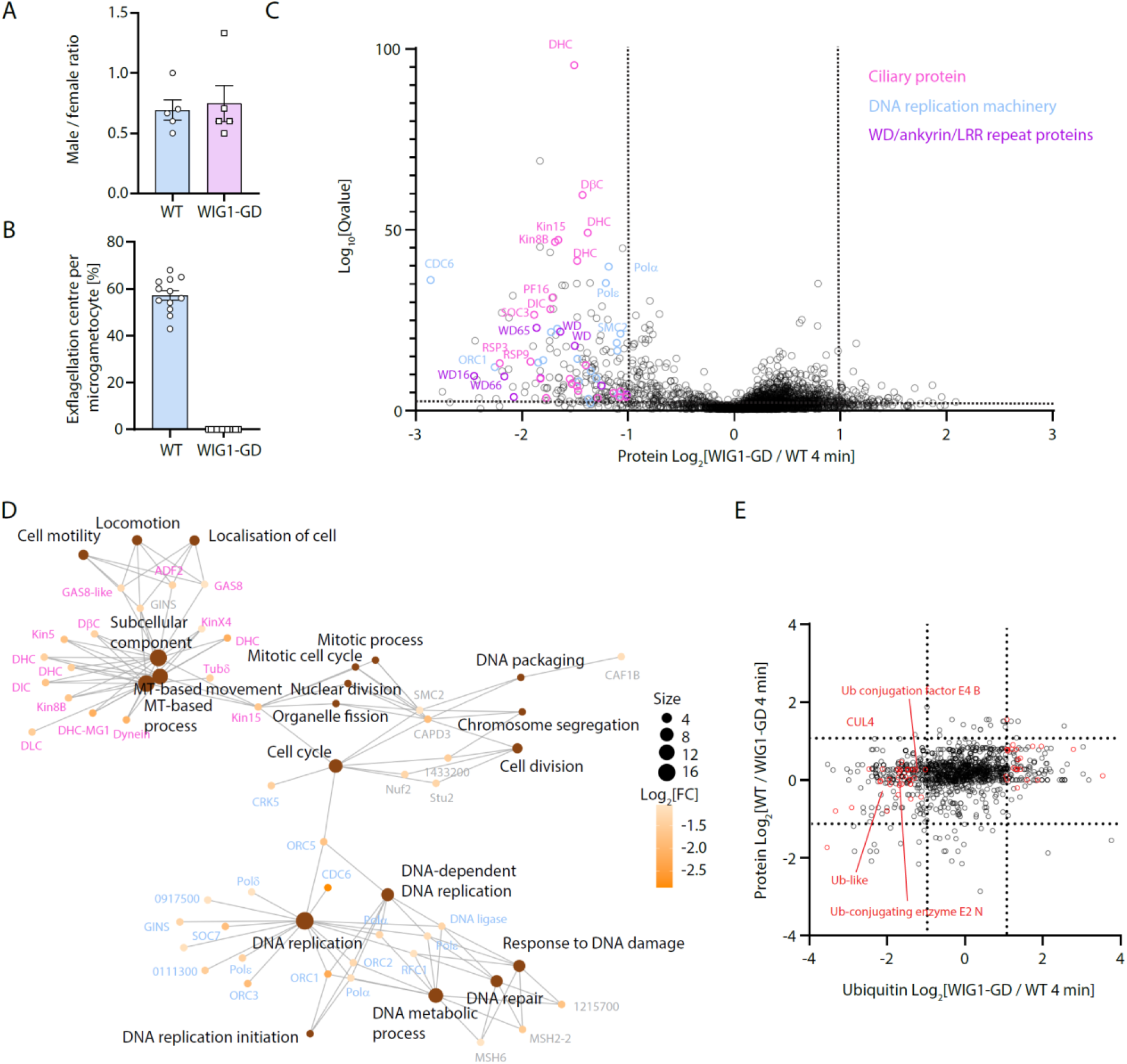
Disruption of WIG1 alters proteostasis of ciliary proteins and components of the DNA replication machinery in *P. berghei* gametocytes. A. Disruption of *wig1* does not alter gametocyte sex ratio (n = 5; two-tailed t-test). **B.** Effect of *wig1* disruption in gametocytes on exflagellation (error bars show standard deviation from the mean; quadruplicates from 3 independent infections). **C.** Volcano plot showing differentially detected proteins in WT and WIG1-GD gametocytes 4 min post-activation (technical duplicates from 2 biological replicates s). **D.** GO term enrichment analysis of proteins down-regulated in 4 min activated WIG1-GD gametocytes. **E.** Plot indicating relative abundance of proteins and corresponding ubiquitinated peptides (technical duplicates from 2 biological replicates) in FBXO1-GD compared to WT gametocytes 4 min post-activation.

## Disruption of *WIG1* alters proteostasis of ciliary proteins and components of the DNA replication machinery

We then set out to exploit the high synchronicity of developing gametocytes to identify whether the defect in exflagellation of WIG1-GD microgametocytes were linked to defective proteostasis or ubiquitination by comparing their proteome and ubiquitome with those of WT parasites (Tables S3 and S4). To do so, we applied an ubiquitin remnant immunoaffinity profiling technique [27]. Trypsin digestion of proteins results in the cleavage at arginine and lysine residues, where the C-terminal Gly-Gly dipeptide of ubiquitin is still attached to the ubiquitinated lysine residue of substrate proteins. Using commercial antibodies that recognize this Gly-Gly remnant, peptides were enriched prior to LC-MS/MS. Data-independent acquisition (DIA)-MS method with a neural network-based data processing were combined to quantify the eluted and flow through fractions. We focused our analysis on activated gametocytes four minutes following stimulation by XA as we previously observed a global rise in ubiquitination at this developmental stage using antibodies targeting ubiquitin [19].

Analysis of the flow through fraction indicated significant differences between the WIG1-GD and WT proteomes with 179 proteins showing at least a two-fold reduction (Q-value < 10^-3^), while only 35 proteins were significantly more detected with the same criteria (Fig. 2C). GO term enrichment analysis indicated that multiple biological processes specific to the microgametocyte were significantly down-regulated. This included microtubule-based movement, mitotic cell cycle and DNA replication (Fig. 2D). Interestingly, proteins associated with microtubule-based movement corresponded to predicted components of the axonemes that mostly correspond to ancestral ciliary proteins [28] including eleven dynein light or heavy chains, kinesins 4X, 5, 8B and 13, SOC3, the armadillo repeat protein PF16, and all three annotated radial spoke proteins. Proteins associated with DNA replication included ORC1 to 5, three subunits of the condesin-2 complex, various subunits of DNA polymerase α, δ, or ε and the cyclin-related kinase 5 (CRK5) that was previously shown to coordinate mitosis and DNA replication during male gametogenesis [10]. In addition, eight proteins containing either WD, LRR or ankyrin repeats were also less detected in the WIG1-GD proteome (Fig. 2C).

In the Gly-Gly enriched fraction, we identified 1,323 ubiquitinated lysine residues mapping on 531 proteins (Table S4). Only 22 proteins with at least one site differentially ubiquitinated were observed. This included CUL4, the ubiquitin-conjugating enzyme E2 N, ubiquitin conjugation factor E4 B and an ubiquitin-like protein suggesting a possible feedback loop (Fig. 2E). However, this dataset did not reveal obvious differences possibly explaining the differential proteome nor the phenotype of WIG1-GD microgametocytes. This is likely due to an earlier requirement for WIG1 during microgametocytogenesis.

## WIG1 is required for efficient DNA replication and formation of functional MTOCs coordinating mitotic spindle and axoneme assembly

In light of the differences observed in the WIG1-GD line at the proteome level, we decided to further refine the phenotype of transgenic parasites during microgametogenesis. Conventional immunofluorescence assays with DNA and α/β-Tubulin staining on 2 min and 14 min activated microgametocytes suggested normal DNA replication with most cells showing expanded DNA staining upon activation. However, staining for α/β-Tubulin did not allow to observe mitotic spindle 2 min post activation and was less homogenous 14 min post-activation suggesting defects in axoneme assembly (Fig. 3A).

**Figure 3.**
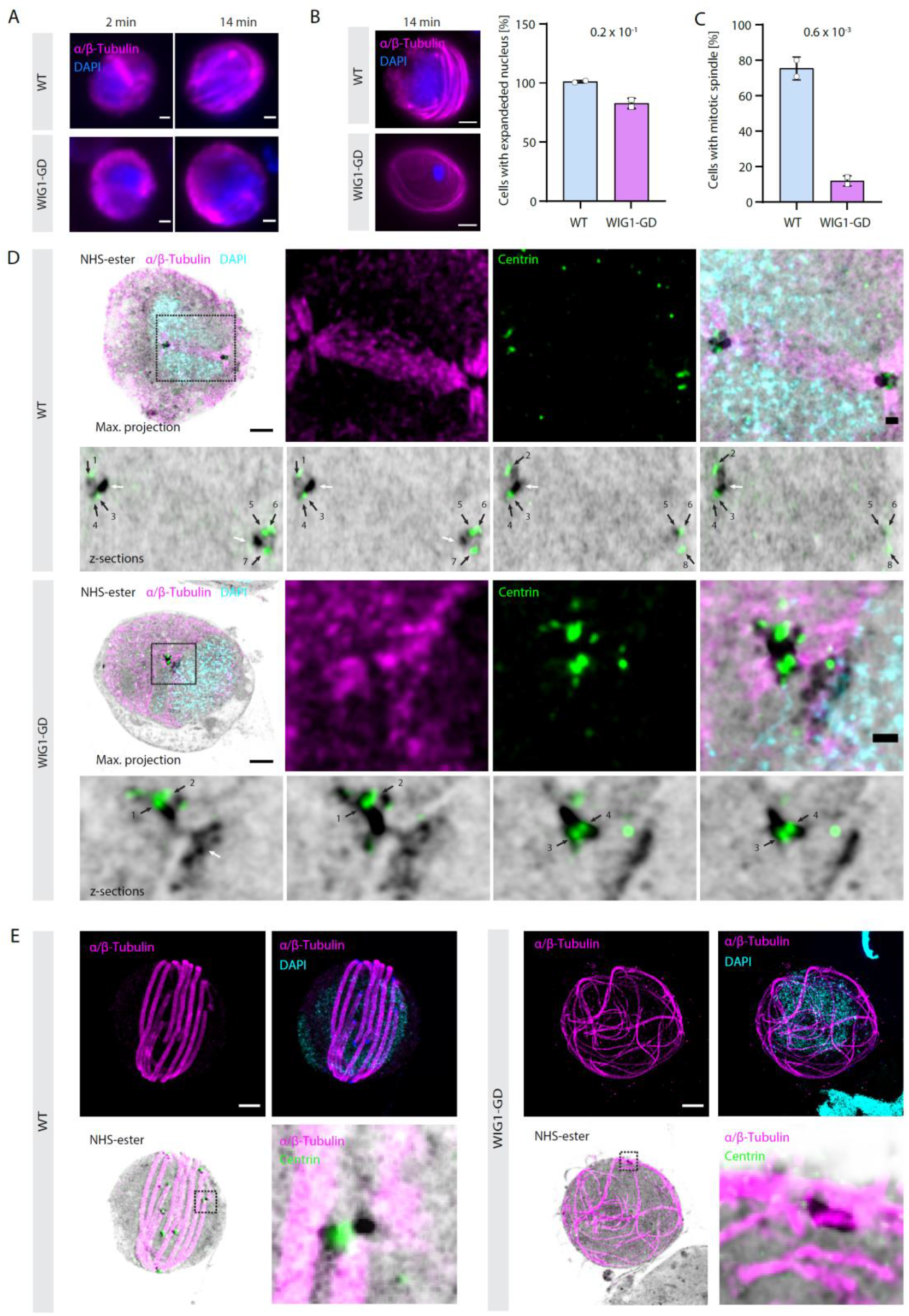
WIG1 is critical for axoneme and mitotic spindle formation and important for DNA replication during *P. berghei* microgametogenesis. A. Widefield images of WT and WIG1-GD microgametocytes 2 min and 14 min post-activation by XA. α/β-Tubulin are shown in magenta and DAPI-stained DNA in blue. Scale bars = 1 µm. **B.** Analysis of DNA replication by U-ExM in *P. berghei* gametocytes. Images on the left show representative images of an expanded nucleus (top) or non-expanded nucleus (bottom). α/β-Tubulin are shown in magenta and DAPI-stained DNA in blue. Scale bars = 5 µm; 148 WT and 169 WIG1-GD gametocytes were analysed from two biological replicates; error bars show standard deviation from the mean; two-tailed t-test. **C.** Analysis of mitotic spindle formation by U-ExM as shown in D in *P. berghei* gametocytes Twenty-five WT and 30 WIG1-GD gametocytes from two biological replicates were analysed, error bars show standard deviation from the mean; two-tailed t-test. **D.** Representative U-ExM images of WT and WIG1-GD gametocytes 1 min post-activation by XA. For each line, top left images are maximum projections of entire cells. Adjacent to each are details of the mitotic spindle (WT) or the related area (WIG1-GD) corresponding to the boxed areas. Below are four confocal sections detailing the MTOC. White arrows: intranuclear body or spindle pole; black arrows: centrin-positive basal bodies. Centrin: green; protein density as assessed by NHS-ester: shades of grey; α/β-Tubulin: magenta; DNA stained with DAPI: cyan. Scale bars entire cells = 5 µm, scale bars insets = 1 µm. **E.** Representative U-ExM maximum projections of WT and WIG1-GD gametocytes 10 min post-activation by XA. Bottom right panels show details of the MTOC corresponding to the boxed areas. Centrin: green; protein density as assessed by NHS-ester: shades of grey; α/β-Tubulin: magenta; DNA stained with DAPI: cyan. Scale bars = 5 µm.

To further investigate potential defects during microgametogenesis, we compared mutant and WT gametocytes with U-ExM [29]. The extent of DAPI staining allowed us to identify microgametocytes that replicated DNA, at least partially, as shown in Fig. 3B. While most WT microgametocytes showed a large DAPI stained nucleus, 19% of WD1-GD microgametocytes displayed a condensed DAPI signal suggestive of impaired DNA replication in these cells [30]. Co-staining of α/β-Tubulin, centrin and NHS-ester of 1 min activated microgametocytes highlighted the absence of mitotic spindle linked to a defect in the replication of the bipartite MTOC (Fig. 3C and D). At this time point, WT microgametocytes displayed two tetrads of four basal bodies at each extremity of the mitotic spindle. This was highlighted by the localisation of centrin dots, as a basal body marker, localising in the NHS-ester dense MTOC. However, in WIG1-GD microgametocytes, most cells displayed a single NHS-ester dense MTOC and a mitotic spindle could be observed in only 15% of observed cells. In addition, no well-defined centrin dots were observed at the base of growing microtubules suggesting defects in the de novo assembly of the basal body and possibly of the intranuclear MTOC (Fig. 3D). In 10 min activated WIG1-GD microgametocytes, no mitotic spindle was visible and only one MTOC could be distinguished, as revealed by NHS-ester staining. Interestingly, while microtubules stained by α/β-Tubulin antibodies nucleated from this single MTOC, they rarely assembled into axonemes as observed in WT microgametocytes (Fig. 3E and Fig. S3).

## Discussion

We have identified and characterised a cullin ring ligase complex that is expressed at multiple stages of the *Plasmodium* lifecycle including asexual blood stages and microgametocytes. This complex is related to the eukaryotic cullin-4 based CRL4 ligases, which have been shown to play various roles including DNA repair, chromatin remodelling, and cell cycle in a wide range of eukaryote including metazoan, yeast and plants [31]. *CUL4* is present as a single gene in *Schizosaccharomyces pombe*, *C. elegans*, *Drosophila*, and *Arabidopsis* whereas mammalian cells express two closely related paralogues, *CUL4A* and *CUL4B*. Here we show that *Plasmodium* CUL4 interacts with RBX1 and DDB1. Through DDB1, these complexes can associate with numerous DDB1– and CUL4-associated factors (DCAF) proteins, which directly interact with specific targets promoting their ubiquitination and subsequent degradation by the proteasome. A characteristic feature of the majority of DCAF proteins that associate with DDB1 is the presence of a WD motif. The WD repeat (also WD40 or beta-transducin repeat) is a short motif of approximately 40 amino acids frequently terminating with a tryptophan-aspartic acid dipeptide. WD-repeat proteins coordinate multi-protein complex assemblies, where the repeating units serve as a rigid scaffold while sequences outside the repeats themselves determine the specificity of the interaction. Immunoprecipitation of CUL4-HA in *P. falciparum* schizonts identified three WD-repeat proteins with a significant enrichment for WIG1 and PF3D7_1410300, while immunoprecipitations of CUL4-HA or DDB1-HA only identified WIG1 in *P. berghei* gametocytes. Genome mining previously identified 80 genes coding for WD40 repeat containing proteins in *Plasmodium* [32] and it is thus possible that many other WD proteins may serve as substrate receptors at different stages of the *Plasmodium* lifecycle.

Depletion of CUL4 in *P. falciparum* strongly impaired intraerythrocytic development of both asexual and sexual stages [25]. This was associated with the deregulation of various pathways at the transcriptome level likely reflecting a pleiotropic requirement for CRL4-based complexes in *Plasmodium*. As depletion of CUL4 likely affects all CRL4s, we wondered whether deletion of a single substrate receptor would allow to dissect requirements for *Plasmodium* CRL4s in more detail. We turned our attention to WIG1 that was detected to interact with CUL4 both in *P. falciparum* schizonts and *P. berghei* gametocytes. To better understand the role of CUL4^WIG1^, we generated a *P. berghei* transgenic clone in which *wig1* was disrupted. This did not affect growth of asexual blood stages suggesting that WIG1 is either not required or that other WD repeat proteins may complement for WIG1 disruption. However, *WIG1* disruption completely abolished formation of microgametes. This defect is likely due to the altered de novo formation of the bipartite MTOCs that coordinate mitosis with axoneme formation during microgametogenesis. This leads to strong defects in the formation of mitotic spindles in the nucleus and the defective axoneme bundling despite nucleation of axonemal microtubules.

Defects observed during microgametogenesis are likely the consequence of altered proteostasis during gametocytogenesis as *WIG1* disruption is associated with down regulation of a large set of ciliary proteins. This includes dynein heavy and light chains, kinesins and radial spoke proteins. Interestingly, previous works showed that deletion of kinesin-8B or radial spoke protein 9 (RSP9) are not required for microtubule nucleation but are necessary for the 9+2 symmetry axoneme formation and hence, production of viable male gametes [33–35]. However, these mutations did not affect mitotic spindle formation suggesting that the observed WIG1-GD phenotype is not solely linked to depletion of Kin-8B or RSP9. Interestingly, deletion of the gene coding for the Serine/arginine-Rich Splicing Factor (SRSF) protein kinase-1 (SRPK1 – PBANKA_0401100) was linked with misincorporation of centrin in the amorphous MTOC leading to similar defects in axoneme and mitotic spindle assembly [29]. Peptides of SRPK1 were identified in PfCUL4-HA and PbCUL4-HA raising the possibility of a functional link between SRPK1 and CRL4^WIG1^. In addition to ciliary proteins, another large set of components of the DNA replication machinery was down-regulated including various subunits of DNA polymerases and most proteins of the origin recognition complex. These changes were also associated with impaired DNA replication defects in at least 19% of observed cells, as assessed by immunofluorescence assays. However, the exact molecular links between CRL4^WIG1^-dependent ubiquitination and the observed phenotypes remain unknown and more work focusing on ubiquitination during microgametocytogenesis will be required to identify the direct targets of CRL4^WIG1^ leading to defective microgametogenesis. We have previously shown an increase in ubiquitination during gametogenesis. As disruption of WIG1 seems to affect gametocyte development it remains unknown whether the protein is active and required during gametogenesis or during later lifecycle stages and further studies will be required to determine the respective role of ubiquitination by APC/C, CRL1/SCF^FBXO1^ or CRL4 ubiquitin ligases during gametogenesis.

Altogether, we show that *Plasmodium* parasites express a CRL4-like complex both in the schizont and gametocyte stages. A putative CRL4 substrate receptor, WIG1, is expressed at both stages. While WIG1 is not essential for asexual blood stages, it is required for DNA replication and essential for the de novo formation of the MTOCs that coordinate mitotic spindle assembly and axoneme biogenesis during microgametogenesis. In addition to the previously described requirement of at least two other ubiquitin E3 ligases, the APC/C [21, 22] and SCF^FBXO1^ [19] complexes, this work further highlights the critical role of protein ubiquitination in the formation of microgametes that are critical for the transmission of malaria.

## Material and methods

### Ethics statement

All animal experiments were conducted with the authorisation numbers GE102 and GE-58-19, according to the guidelines and regulations issued by the Swiss Federal Veterinary Office.

### *P. falciparum* maintenance and transfection

*P. falciparum* strain NF54 was cultured essentially as described in [36]. Briefly, parasites were cultured within O+ human erythrocytes at 5% haematocrit in RPMI 1640 complete medium containing L-glutamine and 25 mM HEPES (Sigma), supplemented with 0.2% sodium bicarbonate, 0.5% Albumax II (Gibco), 0.36 mM hypoxanthine (Sigma) and 50 mg/L gentamicin sulfate (Melford) under a 90% N_2_/5% O_2_/5% CO2 gaseous atmosphere at 37°C, within a hypoxic incubator chamber (Billups-Rothenburg). Parasites were maintained at a parasitaemia of 0.2-10%, calculated by counting the percentage of parasitized erythrocytes of Hemacolor (Sigma) stained thin blood smears using light microscopy. A two-stage process of Percoll separation and selective sorbitol lysis was used to synchronise *Plasmodium* within a 2-hour window. Percoll is a density separation reagent, and sorbitol is an osmotic lysis agent that does not affect ring stage parasites. A 65% Percoll solution was prepared with PBS. *Plasmodium*-infected erythrocyte culture was centrifuged at 800 g for 5 min and the culture supernatant discarded. The cell pellet was resuspended to 10% haematocrit and applied slowly atop the Percoll solution. The volume of the Percoll solution was proportional to the volume of culture to be applied on top (i.e. an 8 mL solution was used to separate a 10 mL 5% haematocrit culture). The layered suspension was centrifuged at 1,500 g for 10 minutes without brake; the parasites migrate to the interface between the 65% Percoll and their culture medium while uninfected RBCs and ring stage infected RBCs are pelleted. The parasites at the interface were recovered and transferred to a new vessel and washed twice in incomplete media before resuspension to 5% haematocrit in complete media. This suspension was agitated at 37°C for 2 hours to facilitate merozoite infection. Following incubation, the culture was pelleted and resuspended in 5% sorbitol (in PBS) for 10 minutes prior to two washes in incomplete media before the infected blood was resuspended and cultured as normal.

Infected erythrocytes at a parasitaemia of 8-10% were collected by centrifugation at 800 g for 10 minutes. The supernatant was aspirated, and the pellet washed in 10 volumes of PBS followed by centrifugation at 800 g for 10 minutes at 4°C. A volume of 0.2% (w/v) saponin equivalent to the starting culture was used to resuspend the cell pellet. This suspension was incubated for 10 minutes at 4°C. Following centrifugation at 3,200 g, the supernatant was removed, and the parasite pellet was washed three times in ice cold PBS. Following the final wash, the supernatant was aspirated, and the pellet flash frozen in liquid nitrogen and stored at –80°C until use.

For transfection, infected human erythrocytes at 10% ring stage parasitaemia were pelleted by centrifugation at 800 g for 5 minutes. The media was aspirated, and 100 µL of the erythrocyte pellet was resuspended in warm 300 µL cytomix (120 mM KCl, 0.15 mM CaCl_2_, 2 mM EGTA, 5 mM MgCl_2_, 10 mM K_2_HPO_4_/KH_2_PO_4_, 25 mM HEPESs pH 7.6) containing 100 ug plasmid DNA. The suspension was electroporated (310 V, 950 µF, 200 Ω, exponential) in a 2mm electroporation cuvette with a GenePulser II (BioRad). Electroporated parasites were immediately transferred into 5 mL complete culture containing 100 µL fresh, washed erythrocytes (50% haematocrit). After two hours the complete media was replaced; after six more hours selection compound was added at an appropriate concentration. Upon resurgence of drug-resistant parasites, cultures were maintained under selection.

*Plasmodium* parasites transfected with the pSLI plasmid [37] containing a 1kb homology region, were treated with 5 nM WR99210 6 hours after transfection. Every day for 5 days the media and WR99210 were refreshed, after this media changes occurred every second day. Following resurgence, parasites were expanded, and the continuing culture was subject to selection with 0.4 mg/mL G418 in complete media. Following resurgence after this selection, parasites were expanded; those parasites being maintained in culture were subjected to double selections at the above concentrations during continuous culture.

### *P. berghei* maintenance and transfection

Parasites were grown and transfected as previously described in [10, 38]. *P. berghei* ANKA strain clone 2.34 together with derived transgenic lines, were grown and maintained in CD1 outbred mice. Six to twelve-week-old mice were obtained from Charles River laboratories, and females were used for all experiments. Mice were specific pathogen free (including *Mycoplasma pulmonis*) and subjected to regular pathogen monitoring by sentinel screening. They were housed in individually ventilated cages furnished with a cardboard mouse house, tunnel and Nestlet, maintained at 21 ± 2°C under a 12 hours light/dark cycle, and given commercially prepared autoclaved dry rodent diet and water *ad libitum*. The parasitaemia of infected animals was determined by microscopy of methanol-fixed and Giemsa-stained thin blood smears.

For gametocyte production, parasites were grown in mice that had been phenyl hydrazine-treated three days before infection. One day after infection, sulfadiazine (20 mg/L) was added in the drinking water to eliminate asexually replicating parasites. Microgametocyte exflagellation was measured three or four days after infection by adding 4 µl of blood from a superficial tail vein to 70 µl exflagellation medium (RPMI 1640 containing 25 mM HEPES, 4 mM sodium bicarbonate, 5% fetal calf serum (FCS), 100 µM xanthurenic acid, pH 7.4) in duplicates at least. An exflagellation event was defined as moving flagellated parasites forming clumps (exflagellation centres) with nearby red blood cells. To calculate the number of exflagellation centres per 100 microgametocytes, the percentage of red blood cells (RBCs) infected with microgametocytes was assessed on Giemsa-stained smears. For gametocyte purification, parasites were harvested in suspended animation medium (SA; RPMI 1640 containing 25 mM HEPES, 5% FCS, 4 mM sodium bicarbonate, pH 7.20) and separated from uninfected erythrocytes on a Histodenz cushion made from 48% of a Histodenz stock (27.6% [w/v] Histodenz [Sigma/Alere Technologies] in 5.0 mM TrisHCl, 3.0 mM KCl, 0.3 mM EDTA, pH 7.20) and 52% SA, final pH 7.2. Gametocytes were harvested from the interface. To induce degradation of AID/H-tagged proteins, 1 mM auxin dissolved in ethanol (0.2% final concentration) was added to purified gametocytes for one hour prior to activation by XA.

Schizonts for transfection were purified from overnight *in vitro* culture on a Histodenz cushion made from 55% of the Histodenz stock and 45% PBS. Parasites were harvested from the interface and collected by centrifugation at 500 *g* for 3 min, resuspended in 25 μL Amaxa Basic Parasite Nucleofector solution (Lonza) and added to 10-20 µg DNA dissolved in 10 µl H_2_O. Cells were electroporated using the FI-115 program of the Amaxa Nucleofector 4D. Transfected parasites were resuspended in 200 µl fresh RBCs and injected intraperitoneally into mice. Parasite selection with 0.07 mg/mL pyrimethamine (Sigma) in the drinking water (pH ∼4.5) was initiated one day after infection. Each mutant parasite was genotyped on a single genomic DNA preparation by PCR using three combinations of primers, specific for either the WT or the modified locus on both sides of the targeted region (experimental designs are shown in Fig. S1 and S2). For allelic replacements, sequences were confirmed by Sanger sequencing using the indicated primers. Parasite lines were cloned when indicated.

### Generation of DNA targeting constructs

The oligonucleotides used to generate and genotype the mutant parasite lines are in Table S5 and a summary of background and generated parasite lines can be found in Table S6. Triple HA or KO targeting vectors were generated using phage recombineering in *Escherichia coli* TSA strain with PlasmoGEM vectors (https://plasmogem.umu.se/pbgem/). For final targeting vectors not available in the PlasmoGEM repository, generation of knock-out and tagging constructs were performed using sequential recombineering and gateway steps as previously described [39, 40]. For each gene of interest (*goi*), the Zeocin-resistance/Phe-sensitivity cassette was introduced using oligonucleotides *goi* HA-F x *goi* HA-R and *goi* KO-F x *goi* KO-R for 3xHA, AID/HA tagging and KO targeting vectors, respectively. Insertion of the GW cassette following gateway reaction was confirmed using primer pairs GW1 x *goi* QCR1 and GW2 x *goi* QCR2. The modified library inserts were then released from the plasmid backbone using NotI. The GD and triple HA targeting vectors were transfected into the 2.34 line. Schematic representations of the targeting constructs as well as WT and recombined loci are shown in Fig. S1 and S2.

## Immunofluorescence assays

*P. berghei* immunofluorescence assays were performed as described in reference [29]. 4% formaldehyde fixed cells were sedimented onto poly-D-Lysine coated microscopy glass slides and briefly incubated with 0.2% TritonX-100 for 5 minutes and blocked with 3% bovine serum albumin for one hour. After primary and secondary antibody incubations and washings, slides were then left to dry and cells were immediately mounted in Vectashield antifade mounting media with DAPI. Images were acquired on a widefield microscope Zeiss Axio Imager M2 using 100X objective.

For *P. falciparum* IFAs, 1 mL of culture was pelleted by centrifugation, and washed once with PBS. The cell pellet was resuspended in 4% formaldehyde (in PBS) and fixed for 1 hour at room temperature with rotation. Fixed cells were pelleted and resuspended in 1 mL 0.1% Triton-X100 (in PBS) for 10 minutes with rotation. Permeabilised cells were pelleted and resuspended in 1% BSA (in PBS) for 1 hour with rotation. A 1:1000 dilution of primary antibody (anti-HA clone 3F10 high affinity, Roche) in 1% BSA at was incubated with the cell suspension for 1 hour at room temperature with rotation. Cells were pelleted and washed 3 times with PBS prior to resuspension in a 1:5000 dilution of secondary antibody (goat anti-rat-AlexaFluor488, ThermoFisher) in 1% BSA. Cells were incubated with rotation for 1 hour at room temperature with rotation. Cells were pelleted and washed 2 times with PBS, then incubated with 1 µg/mL DAPI (in PBS) for 10 minutes. The cells were washed twice with distilled water, before being resuspended to 50% haematocrit in distilled water. A thin smear was prepared on poly-lysine coated microscopy slides and left to air dry before mounting in ProLong Diamond antifade mountant (ThermoFisher). Images were captured using a fluorescence widefield microscope and processed using Fiji/Image J.

### Expansion microscopy

U-ExM was performed as described in [29, 41]. Formaldehyde fixed samples were settled on a 12 mm round Poly-D-Lysine (A3890401, Gibco) coated coverslips for 10-12 minutes. Coverslips were incubated for five hours in 1.4% formaldehyde (FA)/ 2% acrylamide (AA) at 37°C. Thereafter gelation was performed in ammonium persulfate (APS)/Temed (10% each)/Monomer solution (23% Sodium Acrylate; 10% AA; 0,1% BIS-AA in PBS) for 1 hour at 37°C. Gels were denatured for 1 hour and 30 minutes at 95°C. After denaturation, gels were incubated in distiller water overnight for complete expansion. The following day, gels were washed in PBS twice for 15 minutes to remove excess of water. Gels were then incubated with primary antibodies at 37°C for 3 hours, and washed three times for ten minutes in PBS-Tween 0.1%. Incubation with secondary antibodies was performed for three hours at 37°C followed by three washes of ten minutes each in PBS-Tween 0.1% (all antibody incubation steps were performed with 120–160 rpm shaking at 37°C). Directly after antibody staining, gels were incubated in 1 ml of 594 NHS-ester (Merck: 08741) diluted at 10 μg/mL) in PBS for one hour and 30 minutes at room temperature on a shaker. The gels were then washed three times for 15 minutes with PBS-Tween 0.1% and expanded overnight in ultrapure water. Gel pieces of 1cm x 1cm were cut from the expanded gel and attached on 24 mm round Poly-D-Lysine (A3890401, Gibco) coated coverslips to prevent gel from sliding and to avoid drifting while imaging. The coverslip was then mounted on a metallic O-ring 35mm imaging chamber (Okolab, RA-35-18 2000–06) and imaged. Images were acquired on Leica TCS SP8 microscope with HC PL Apo 100x/ 1.40 oil immersion objective in lightning mode to generate deconvolved images. System optimised z stacks were captured between frames using HyD as detector. Images were processed with ImageJ and LAS X softwares. A list of used antibodies and dyes can be found in Table S6.

## Immunoprecipitations

### Sample preparation for proteomic characterisation of *P. falciparum* schizonts

For PfCUL4-HA immunoprecipitations, 490 mL late stage schizont culture at 8 % parasitaemia was grown and harvested per sample. Frozen parasite pellets were thawed on ice, before being resuspended in ice cold, sterile filtered lysis buffer (50 mM HEPES pH 7.5, 150 mM NaCl, 1% Triton-X100, Complete Mini protease inhibitor). The cell suspension was lysed by 5 freeze-thaw cycles with agitation and vortex mixing between each cycle. The insoluble fraction was pelleted by centrifugation at 17,000 g for 30 minutes at 4°C, and the soluble fraction aspirated. Protein concentration in the soluble fraction was estimated using Pierce BCA protein assay kit (ThermoFisher) following the manufacturer’s instructions. Total protein concentration was normalised to 10 mg input across samples. 25 μL pre-washed magnetic anti-HA resin (ThermoFisher) was added to each lysate sample and incubated for 2 hours at 4°C with rotation. Beads were gathered by magnet, the supernatant aspirated, and the beads washed 3 times (50 mM HEPES pH 7.5, 150 mM NaCl, 0.1% Triton-X100, Complete Mini protease inhibitor) with a change of vessel between the second and third wash. The wash solution was aspirated, and the beads were resuspended in 25 μL of 1 mg/mL HA peptide (in 15 mM Tris pH 7.4, 150 mM NaCl, 0.1% SDS, 0.5% NP40) and incubated for 30 minutes at 37°C with agitation. The eluant was collected and introduced into an SDS-PAGE gel for 5 minutes. The gel was fixed (as above) and transferred to the mass spectrometry facility for processing and LC-MS/MS (Department of Biochemistry, University of Cambridge).

For analysis of the HA-mediated immunoprecipitation of PfCUL4-HA, raw data was provided as output from the mass spectrometry facility at the University of Cambridge. This was used as input for MaxQuant v.2.0.3.1. Parameters were set as default with exceptions: deamidation (NQ) and GlyGly (K) were included as variable modifications; label min ratio count was set to 1 and deamidation was included as a modification used in protein quantification, while discard unmodified counterpart peptides was unchecked; the FTMS MS/MS was set to 0.05 Da and the ITMS MS/MS match tolerance was set to 0.6 Da; and iBAQ LFQ was checked. The PlasmoDB v61 proteome fasta file was used as the reference proteome file. The resultant ProteinGroup output file was input into Perseus v1.6.15.0 for further analysis using the iBAQ LFQ values. Column values were filtered based on categorical column: only identified by site, reverse, and potential contaminant. Samples were grouped as control or experimental by categorical row annotation. iBAQ LFQ values were transformed by log2(x), and invalid values filtered with a minimum requirement of 2 valid values per group in the case triplicates. Invalid values were imputed from a normal distribution: width (0.3), down shift (1.8). As a triplicate experiment, a volcano plot was generated using 250 randomisations, FDR 0.05, and S0 0.1; enriched proteins positioned above the p<0.05 threshold were taken as hits.

### Sample preparation for proteomic characterisation of *P. berghei* gametocyte immunoprecipitates

Co-immunoprecipitations (IPs) of proteins were performed with purified gametocytes from two or three independent biological replicates, as previously described [10, 19, 38]. Samples were fixed for 10 minutes with 1% formaldehyde. Parasites were lysed in RIPA buffer (50 mM Tris HCl pH 8, 150 mM NaCl, 1% NP-40, 0.5% sodium deoxycholate, 0.1% SDS) and the supernatant was subjected to affinity purification with anti-HA antibody (Sigma) conjugated to magnetics beads. Beads were re-suspended in 100 μl of 6 M urea in 50 mM ammonium bicarbonate (AB). Two μl of 50 mM dithioerythritol (DTE) were added and the reduction was carried out at 37°C for 1h. Alkylation was performed by adding 2 μl of 400 mM iodoacetamide for 1 h at room temperature in the dark. Urea was reduced to 1 M by addition of 500 μl AB and overnight digestion was performed at 37 °C with 5 μl of freshly prepared 0.2 μg/μl trypsin (Promega) in AB. Supernatants were collected and completely dried under speed-vacuum. Samples were then desalted with a C18 microspin column (Harvard Apparatus) according to manufacturer’s instructions, completely dried under speed-vacuum and stored at –20°C.

### Liquid chromatography electrospray ionisation tandem mass spectrometry (LC-ESI-MSMS)

Samples were diluted in 20 μl loading buffer (5% acetonitrile [CH_3_CN], 0.1% formic acid [FA]) and 2 μl were injected onto the column. LC-ESI-MS/MS was performed either on a Q-Exactive Plus Hybrid Quadrupole-Orbitrap Mass Spectrometer (Thermo Fisher Scientific) equipped with an Easy nLC 1000 liquid chromatography system (Thermo Fisher Scientific) or an Orbitrap Fusion Lumos Tribrid mass Spectrometer (Thermo Fisher Scientific) equipped with an Easy nLC 1200 liquid chromatography system (Thermo Fisher Scientific). Peptides were trapped on an Acclaim pepmap100, 3 μm C18, 75 μm x 20 mm nano trap-column (Thermo Fisher Scientific) and separated on a 75 μm x 250 mm (Q-Exactive) or 500 mm (Orbitrap Fusion Lumos), 2 μm C18, 100 Å Easy-Spray column (Thermo Fisher Scientific). The analytical separation used a gradient of H_2_O/0.1% FA (solvent A) and CH_3_CN/0.1 % FA (solvent B). The gradient was run as follows: 0 to 5 min 95 % A and 5 % B, then to 65 % A and 35 % B for 60 min, then to 10 % A and 90 % B for 10 min and finally for 15 min at 10 % A and 90 % B. Flow rate was 250 nL/min for a total run time of 90 min.

Data-dependant analysis (DDA) was performed on the Q-Exactive Plus with MS1 full scan at a resolution of 70,000 Full width at half maximum (FWHM) followed by MS2 scans on up to 15 selected precursors. MS1 was performed with an AGC target of 3 x 10^6^, a maximum injection time of 100 ms and a scan range from 400 to 2000 m/z. MS2 was performed at a resolution of 17,500 FWHM with an automatic gain control (AGC) target at 1 x 10^5^ and a maximum injection time of 50 ms. Isolation window was set at 1.6 m/z and 27% normalised collision energy was used for higher-energy collisional dissociation (HCD). DDA was performed on the Orbitrap Fusion Lumos with MS1 full scan at a resolution of 120,000 FWHM followed by as many subsequent MS2 scans on selected precursors as possible within a 3 sec maximum cycle time. MS1 was performed in the Orbitrap with an AGC target of 4 x 10^5^, a maximum injection time of 50 ms and a scan range from 400 to 2000 m/z. MS2 was performed in the Ion Trap with a rapid scan rate, an AGC target of 1 x 10^4^ and a maximum injection time of 35 ms. Isolation window was set at 1.2 m/z and 30% normalised collision energy was used for HCD.

### Database searches

Peak lists (MGF file format) were generated from raw data using the MS Convert conversion tool from ProteoWizard. The peak list files were searched against the PlasmoDB_*P.berghei* ANKA database (PlasmoDB.org[42], release 38, 5076 entries) combined with an in-house database of common contaminants using Mascot (Matrix Science, London, UK; version 2.5.1). Trypsin was selected as the enzyme, with one potential missed cleavage. Precursor ion tolerance was set to 10 ppm and fragment ion tolerance to 0.02 Da for Q-Exactive Plus data and to 0.6 for Lumos data. Variable amino acid modifications were oxidized methionine and deamination (Asn and Gln) as well as phosphorylated serine, threonine and tyrosine. Fixed amino acid modification was carbamidomethyl cysteine. The Mascot search was validated using Scaffold 4.8.4 (Proteome Software) with 1% of protein false discovery rate (FDR) and at least 2 unique peptides per protein with a 0.1% peptide FDR.

### Principal component analyses (PCA)

Proteins identified in the WT control from reference [10] were removed from the analysis. For PCA analysis, IPs of FBXO1-HA performed under the same experimental conditions [10] were used as an additional control. Quantitative value (Normalized Total Spectra) was calculated by Scaffold. Missing values were imputed with the minimum fixed value of 0.5 and Principal Component Analysis were computed and plotted using R.

## Proteomic and ubiquitinomic analyses

Proteomic and ubiquitinomic analyses were performed as previously described in reference [19].

### Sample preparation for proteomic characterisation of gametocytes

Cell lysis of two independent biological replicates was performed in 100 µl of 2% SDS, 25 mM NaCl, 50 mM Tris (pH 7.4), 2.5 mM EDTA and 20 mM TCEP supplemented with 1x Halt™ protease and phosphatase inhibitor. Samples were vortexed and then heated at 95°C for 10 min with 400 rpm mixing with a thermomixer. DNA was sheared with four sonication pulses of 10 s each at 50% power. Samples were centrifuged for 30 min at 17’000 g and supernatants were collected. A Pierce protein assay was performed for each sample. Samples were then diluted with 200 µl of 50 mM Tris (pH 7.4) and incubated with 48 µl of iodoacetamide 0.5 M for 1 hour at room temperature. Proteins were digested based on the FASP method using Amicon® Ultra-4, 30 kDa as centrifugal filter units (Millipore). Trypsin (Promega) was added at a 1:80 enzyme:protein ratio and digestion was performed overnight at room temperature. The resulting peptide samples were desalted with Pierce^TM^ Peptide Deslting Spin Column (Thermo Fisher Scientific) according to manufacturer’s instruction; peptide concentration was determined using a colorimetric peptide assay (Thermo Fisher Scientific) then completely dried under speed-vacuum.

### Enrichment of ubiquitinated peptides

The PTM-Scan ubiquitin remnant motif (K-ɛ-GG) kit (Cell Signaling Technology, Kit #5562) was used and instructions described by the manufacturer were followed. For each ubiquitin remnant peptide enrichment immunoprecipitation (independent biological duplicates, each split in technical duplicates), approximately 8-10 mg of total protein from purified gametocytes were used. Gametocytes were harvested from about 8-10 ml of *P. berghei* infected blood in suspended animation and treated with 1 µm MG132 for 1 hour at 37C. Ubiquitin remnant peptide enrichment was performed as described in[43] with some modifications. Cell pellets were lysed in 5 mM iodoacetamide (IAA – I1149, Sigma-Aldrich) and 5% SDS/100 mM triethylammonium bicarbonate (TEAB – T7408, Sigma-Aldrich), and processed by ultrasonic probe and heated at 90C for 10 minutes. Samples were then processed again by ultrasonic probe five times 30 seconds on ice. Lysates were cleared by centrifugation at 16,000 g for 15 minutes. Protein concentration was measured with the Pierce 660 nm Protein Assay (Thermo), reduced with 10 mM tris(2-carboxyethyl)phosphine (TCEP – 77720, Thermo) at 56°C for 15 minutes and alkylated by 10 mM IAA at RT for 30 minutes. Proteins were then precipitated by chloroform/methanol. One milliliter of 100 mM TEAB was added to the protein pellet and the mixture was left in ultrasonic bath to disperse the pellet. One hundred micrograms of trypsin (Pierce) was added and incubated at 37C for 2 hours. Another 100 μg of trypsin was added and incubated for further 15 hours. The protein digest was then heated at 70°C for 10 minutes and dried in a SpeedVac. The PTMScan^®^ *IAP buffer* was then added to the pellet and the mix was further sonicated to dissolve the pellet. The sample was then centrifuged at 16,000 g at 4°C to clear the lysate. To the cleared peptide digest, a vial of Cell Signaling Technology antibody-beads were added and incubated with rotation for two hours at room temperature followed by an overnight incubation at 4°C. The mixture was centrifuged at 2000 g for 30 seconds, ubiquitin-peptide enriched beads were washed twice with the IAP buffer and once with ice cold HPLC water. Samples were finally incubated twice with 55 µl of 0.15% TFA for 10 minutes to elute ubiquitin-peptides. One hundred microliters of supernatant sample and 110 µL of IP (GlyGly enriched sample) were desalted with a C18 macrospin column and a C18 microspin column, respectively, (Harvard Apparatus, Holliston, MA, USA) according to manufacturer’s instructions. A PierceTM Colorimetric Peptide assay (cat. No. 23275) was used to quantify the peptide amount after the desalting procedure. The total amount of peptides was of 24 µg and 68 µg in the GlyGly enriched and supernatant samples, respectively.

### ESI-LC-MSMS

Duplicate samples were dissolved at 1 μg/μl with loading buffer (5% CH3CN, 0.1% FA). Biognosys iRT peptides were added to each sample and 2 μg of peptides were injected on column. LC-ESI-MS/MS was performed on an Orbitrap Fusion Lumos Tribrid mass spectrometer (Thermo Fisher Scientific) equipped with an Easy nLC1200 liquid chromatography system (Thermo Fisher Scientific). Peptides were trapped on a Acclaim pepmap100, C18, 3 μm, 75 μm x 20 mm nano trap-column (Thermo Fisher Scientific) and separated on a 75 μm x 500 mm, C18 ReproSil-Pur (Dr. Maisch GmBH), 1.9 μm, 100 Å, home-made column. The analytical separation was run for 135 min using a gradient of H2O/FA 99.9%/0.1% (solvent A) and CH3CN/H2O/FA 80.0%/19.9%/0.1% (solvent B). The gradient was run from 8 % B to 28 % B in 110 min, then to 42% B in 25 min, then to 95%B in 5 min with a final stay of 20 min at 95 % B. Flow rate was of 250 nL/min and total run time was of 160 min. Data-Independent Acquisition (DIA) was performed with MS1 full scan at a resolution of 60,000 (FWHM) followed by 30 DIA MS2 scan with fix windows. MS1 was performed in the Orbitrap with an AGC target of 1 x 106, a maximum injection time of 50 ms and a scan range from 400 to 1240 m/z. DIA MS2 was performed in the Orbitrap using higher-energy collisional dissociation (HCD) at 30%. Isolation windows was set to 28 m/z with an AGC target of 1 x 106 and a maximum injection time of 54 ms.

### Data analysis

DIA raw files for IP samples and Supernatant samples were loaded separately on Spectronaut v.15 (Biognosys) and analysed by directDIA using default settings (two .SNE file have been generated). Briefly, data were searched against *Plasmodium berghei* ANKA database (PlasmoDB.org, release 49, 5076 entries). Trypsin was selected as the enzyme, with two potential missed cleavage. Variable amino acid modifications were oxidized methionine and GlyGly (GG) derivatisation of lysine (+ 114 Da) (K). Fixed amino acid modification was carbamidomethyl cysteine. Both peptide precursor and protein FDR were controlled at 1% (Q value < 0.01). Single Hit Proteins were excluded for Super Natant samples. For quantitation, Top 3 precursor area per peptides were used, “only protein group specific” was selected as proteotypicity filter and normalization was set to “automatic”. A paired t-test was applied for differential abundance testing. The following parameters were used: Quantity at the MS2 level, quantity type = area, data filtering by Qvalue, and cross run normalization selected. Proteins and peptides were considered to have significantly changed in abundance with a Qvalue ≤ 0.05 and an absolute fold change FC≥ |1.5| (log_2_FC ≥ |0.58|).

### Enrichment analysis

Gene Ontologies term enrichment analysis, as well as associated plots, were performed with ClusterProfiler [44, 45] R package, using the EnrichGO function as previously describe in reference [19]. Enrichment tests were calculated for GO terms, based on hypergeometric distribution. P-value cutoff was set to 0.05. The gofilter function was use prior to cnetplot drawing, to filter results at specific levels.

### Softwares

Image J 1.53, and LAS X (Leica version 3.5.7.23225) were used for image analysis. Excel (v1108), GraphPad Prism 9 (9.5.1) were used for data and statistical analysis. Bio-Rad Image Lab (6.1) was used for western blot analysis. The R package was used for proteomic data analysis and representation. ClusterProfiler (3.8) was used for Gene Ontologies term enrichment analysis. Spectronaut v.15 (Biognosys) used for DIA analysis. Mascot (Matrix Science, London, UK; version 2.5.1) was used for the peak list files searches against the PlasmoDB_P.berghei ANKA database (PlasmoDB.org, release 38). ZEN 2.6 (Zeiss) and LAS X (Leica version 3.5.7.23225) were used for image acquisition.

## Acknowledgements

We thank the excellent service at the bioimaging (François Prodon, Nicolas Liaudet and Olivier Brun) and proteomic (Nadia Walter, Domitille Schvartz, Carla Pasquarello and Alexandre Hainard) core facilities at the Faculty of Medicine (University of Geneva). This work was supported by the Swiss National Science Foundation (31003A_179321 and 310030_208151) to MB, a BBSRC Project Grant BB/R001642/1 held by KAT and a Wellcome Trust Doctoral Training Program studentship awarded to CS. *P. falciparum* mass spectrometry was done at the Cambridge Centre for Proteomics Core Facility under the guidance of Dr. Mike Deery whom we thank for his help.

## Data availability

All data needed to evaluate the conclusions in the paper are present in the paper and/or the Supplementary Materials. The WIG1 proteomic and ubiquitinomic data have been deposited to the ProteomeXchange Consortium via the PRIDE partner repository (http://proteomecentral.proteomexchange.org) with the dataset identifier PXD035557.

## Supplementary material

### Supplementary figures

**Figure S1.**
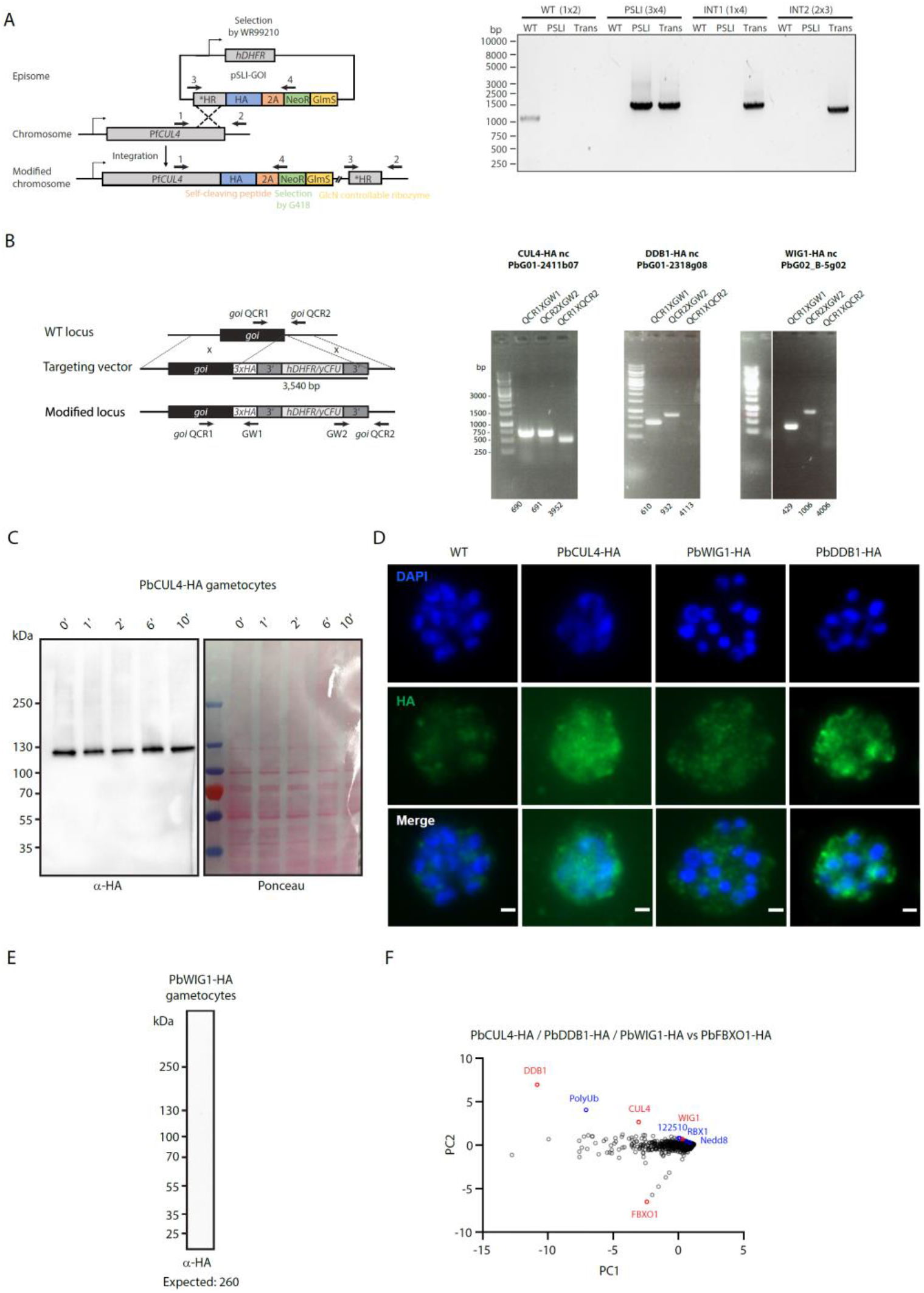
**A.** Genetic modification strategy for PfCUL4 HA tagging and genotyping data. **B.** Genetic modification strategy for HA tagging in *P. berghei* and genotyping data. Oligonucleotides used for PCR genotyping are indicated and agarose gels for corresponding PCR products from genotyping reactions are shown. The same marker size was used for all genotyping gels. PCR product sizes are indicated below each lane; NA: not amplifiable. In the line names nc is for non-clonala, the ID below is the PlasmoGEM vector from which the construct was derived. **C.** Western blot analysis of PbCUL4-HA gametocyte lysates over the course of gametogenesis. The Ponceau staining serves as a loading control. The blot is representative of 2 independent replicates. **D.** Localisation of PbCUL4-HA, PbDDB1-HA and WIG1-HA (all green), by widefield immunofluorescence in segmenting schizonts. DNA is stained with DAPI (blue). Scale bars: schizont = 1 µm. **E.** Western blot analysis of a gametocyte lysate from the line expressing endogenously HA-tagged WIG1 does not allow to detect the fusion protein. The blot is representative of 2 independent replicates. **F.** Quantitative value (Normalized Total Spectra) for proteins co-purifying with PbCUL4-HA, PbDDB1-HA, PbWIG1-HA and PbFBXO1-HA (control) following immunoprecipitation, and displayed in first and second principal components (n = 2 biological replicates for each bait). Red denotes immunoprecipitated proteins and blue possible components of the CRL4 complex.

**Figure S2.**
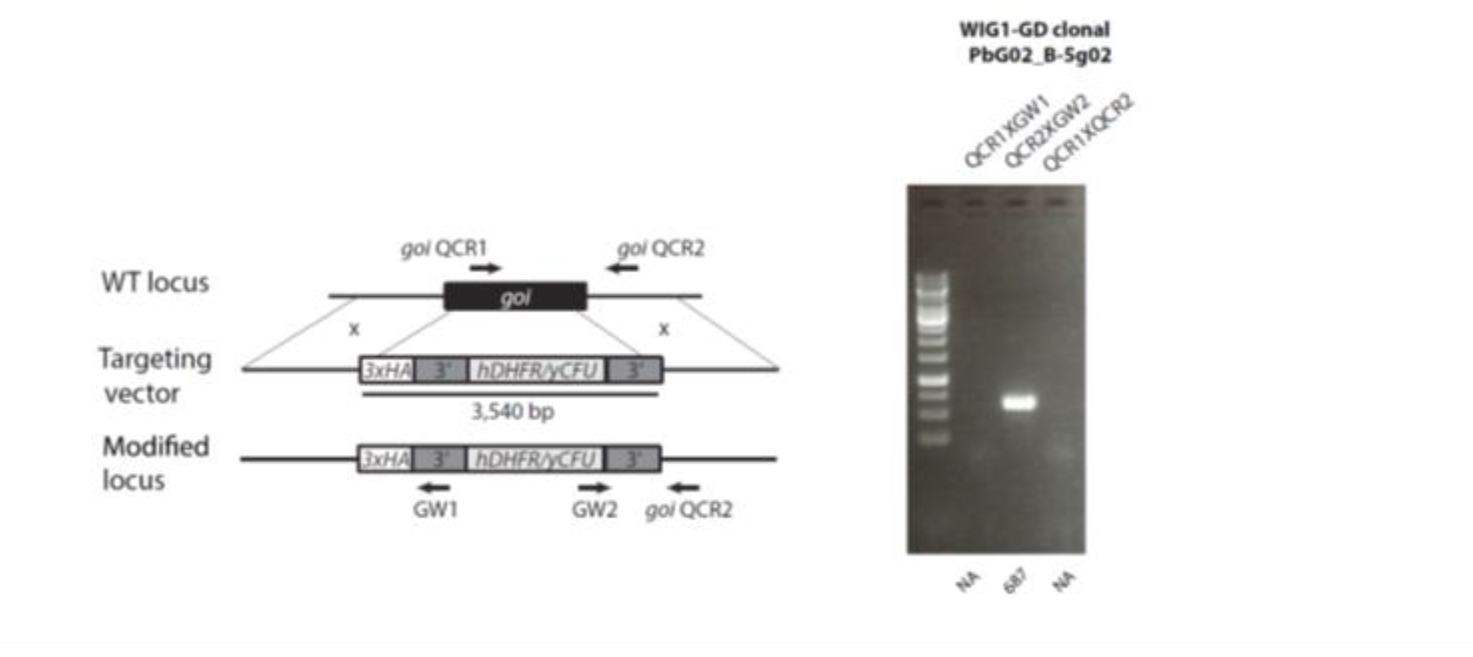
Genetic modification strategy for *WIG1* gene disruption in *P. berghei* and genotyping data. Oligonucleotides used for PCR genotyping are indicated and agarose gels for corresponding PCR products from genotyping reactions are shown. The same marker size was used for all genotyping gels. PCR product sizes are indicated below each lane; NA: not amplifiable. The ID below the line name is the PlasmoGEM vector from which the construct was derived.

**Figure S3.**
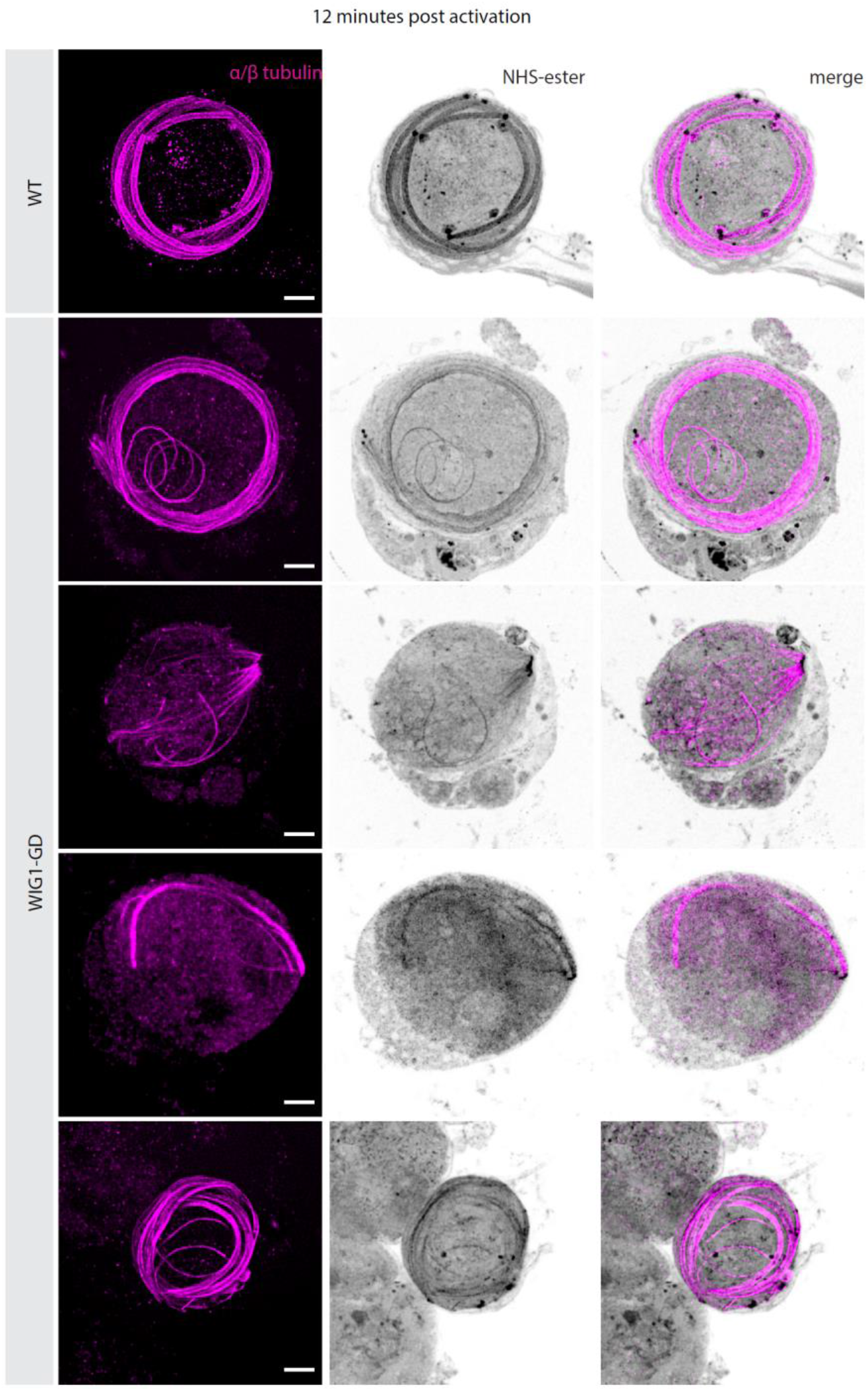
Gallery of maximum projections showing the effect of WIG1 disruption on microgametogenesis as observed by U-ExM 12 minutes after activation by XA. α/β-Tubulin: magenta; amine reactive groups/NHS-ester: shades of grey. Scale bars = 5 µm.

### Supplementary tables

**Table S1.** Proteins detected in CUL4-HA immunoprecipitates from *P. falciparum* schizonts.

**Table S2.** Proteins detected in CUL4-HA, DDB1-HA and WIG1-HA immunoprecipitates from *P. berghei* gametocytes.

**Table S3.** Proteome comparison of WT and WIG1-GD gametocytes four minutes post-activation by XA.

**Table S4.** Ubiquitome comparison of WT and WIG1-GD gametocytes four minutes post-activation by XA.

**Table S5.** Oligonucleotides used in this study.

**Table S6.** Main reagents used or generated in this study.

